# An amino acid-resolution interactome for motile cilia illuminates the structure and function of ciliopathy protein complexes

**DOI:** 10.1101/2023.07.09.548259

**Authors:** Caitlyn L. McCafferty, Ophelia Papoulas, Chanjae Lee, Khanh Huy Bui, David W. Taylor, Edward M. Marcotte, John B. Wallingford

**Author notes:** Correspondence to: C.L.M.,; E.M.M.,; J.B.W. These authors contributed equally (Caitlyn L. McCafferty, Ophelia Papoulas, Chanjae Lee).

## Abstract

Motile cilia are ancient, evolutionarily conserved organelles whose dysfunction underlies motile ciliopathies, a broad class of human diseases. Motile cilia contain myriad different proteins that assemble into an array of distinct machines, so understanding the interactions and functional hierarchies among them presents an important challenge. Here, we defined the protein interactome of motile axonemes using cross-linking mass spectrometry (XL/MS) in *Tetrahymena thermophila*. From over 19,000 XLs, we identified 4,757 unique amino acid interactions among 1,143 distinct proteins, providing both macromolecular and atomic-scale insights into diverse ciliary machines, including the Intraflagellar Transport system, axonemal dynein arms, radial spokes, the 96 nm ruler, and microtubule inner proteins, among others. Guided by this dataset, we used vertebrate multiciliated cells to reveal novel functional interactions among several poorly-defined human ciliopathy proteins. The dataset therefore provides a powerful resource for studying the basic biology of an ancient organelle and the molecular etiology of human genetic disease.

**Highlights:**

- Over 4,700 distinct amino-acid cross-links reveal the composition, structural organization, and conformational dynamics of proteins in motile ciliary axonemes.
- Dense interaction networks are defined for tubulins, Intraflagellar Transport complexes, axonemal dyneins, radial spokes, the CCDC39/40 96nm molecular ruler, and other complexes.
- These data reveal the placement of multiple adenylate kinases in the central apparatus and radial spokes of motile axonemes, a new microtubule-associated protein complex of CFAP58 and CCDC146, and insights into the activity of ciliopathy protein MAATS1/CFAP91.
- The data also provide the first known molecular defect resulting from loss of the human ciliopathy protein ENKUR.

## Introduction

Vertebrate animals require motile cilia to generate fluid flows that are critical to the development and homeostasis of several organ systems. Accordingly, genetic defects that disrupt motile cilia function are associated with a broad class of human diseases known as motile ciliopathies, which involve repeated sinopulmonary disease, bronchiectasis, cardiac defects such as heterotaxy, situs anomalies, and infertility^1,2^. Human genomic studies have implicated many dozens of genes in motile ciliopathy, but that progress has not been matched by a mechanistic understanding of the corresponding proteins’ functions.

This stubborn gap in our understanding results in part from the remarkable complexity of the motile cilia proteome. By most estimates, motile cilia contain roughly 1,000 distinct proteins, and these are assembled into a diverse array of multi-protein machines (**Figure 1A**). Most notable are the axonemal dynein motors that decorate the outside of the microtubule doublets, drive ciliary beating, and tune the ciliary waveform^3,4^. Present in repeated units along the axoneme, these motors comprise at least two subtypes of outer dynein arms (ODAs) and seven sub-types of inner dynein arms (IDAs). In addition to these motors, normal beating in most motile cilia also relies upon the central pair microtubules and their associated central apparatus^5^, the radial spokes that project from the surrounding doublets toward the central pair^6^, and the microtubule inner proteins (MIPs) that nestle within the outer doublet microtubules^7^ (**Figure 1A**). Variants in genes encoding subunits in any one of these systems are sufficient to cause motile ciliopathy^1^.

**Figure 1:**
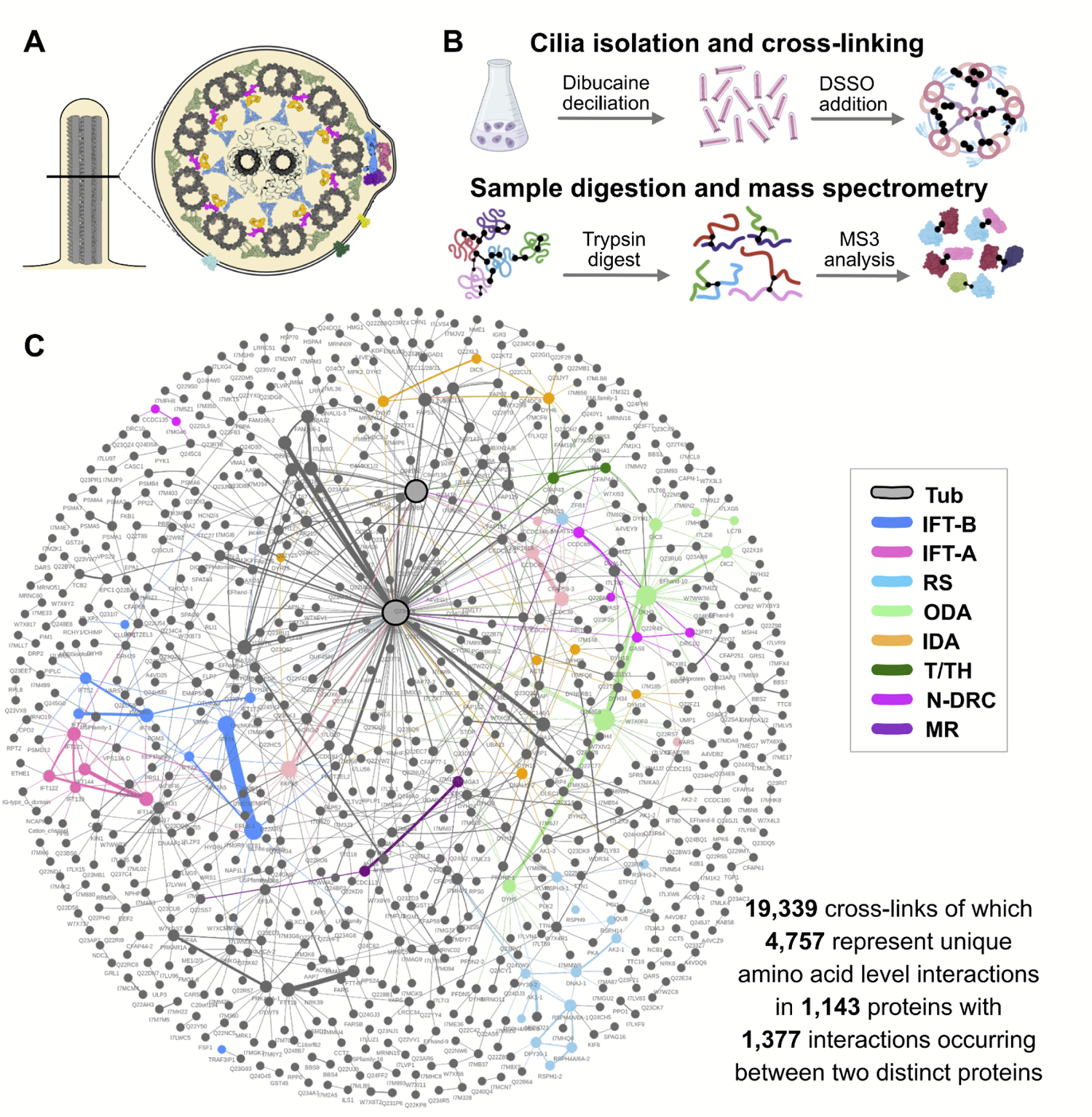
XL/MS reveals the motile ciliary interactome. **(A)** A cross-section of a motile cilium is shown, illustrating key components known from cryo-electron microscopy (cryo-EM) and cryo-electron tomography (cryo-ET) studies, including the 9 microtubule (MT) doublets and central MT pair. Colors match the interaction network below. **(B)** The general protocol used to obtain the motile cilia interactome by cross-linking/mass spectrometry (XL/MS). Cilia were released from *Tetrahymena* using dibucaine, and the membrane permeable cross-linker DSSO was added to the intact cilia to capture native protein interactions. This data was supplemented by XL/MS experiments on native biochemical fractions of disrupted cilia to produce the motile cilia interaction network shown in **(C)**. Colored nodes represent proteins highlighted in **Figure 2**. Tub, tubulin; IFT-B, intraflagellar-B complex; IFT-A, intraflagellar-A complex; RS, radial spoke; ODA, outer dynein arm; IDA, inner dynein arm; T/TH, tether/tether head; N-DRC, nexin-dynein regulatory complex; MR, molecular ruler.

Further complexity arises because each of these machineries displays both radial and proximodistal patterns of asymmetry^8^, and these patterns are established by a host of adaptor proteins. For example, ciliopathy patients with variants in *CCDC39* or *CCDC40* genes display severe derangement of the axoneme and especially poor patient outcomes^9–11^, because the CCDC39 / CCDC40 complex is the “ruler” that sets the length for the repeating units of ODAs and IDAs^12^. By contrast, other adaptor complexes position only specific ODA and IDA sub-types, and loss of any of these results in comparatively milder axonemal phenotypes characterized by loss of only specific motors (e.g. refs. ^13,14^). Finally, for many ciliopathy-associated proteins, no structural or molecular defect in patient axonemes has yet been identified, underscoring the fact that better diagnosis of motile ciliopathies will require a deeper understanding of protein networks in motile axonemes^15–17^.

This still-incomplete picture of the protein interaction landscape in motile cilia reflects a general challenge in biology, and improved protein interactomes are powerful drivers for discovery^18^.

Targeted approaches such as affinity-purification/mass spectroscopy (AP/MS)^19–21^ and proximity-labeling^22,23^ have proven very effective. However, these tag-based methods are challenging to apply systematically to specialized cell types such as those bearing motile cilia, so recent efforts have been made to develop label-free methods for interactome mapping (e.g. refs. ^24–28^). One such method is cross-linking/mass spectrometry (XL/MS), which derivatizes closely positioned reactive amino acids (e.g., lysines), thereby identifying pairs of closely associated proteins^29–32^. By detecting very close amino acid neighbors, XL/MS has the added power of providing structural insights, since the measured cross-links constrain the relative 3D orientation and packing of the interaction partners. Thus, XLs can be combined with protein structure prediction or cryo-electron tomography to build detailed 3D models of protein complexes^33–36^.

Here, we used XL/MS to determine the protein-protein interactome of motile cilia. We generated over 19,000 XLs representing 4,757 unique binary amino acid interactions, providing both broad and deep coverage of the motile ciliary interactome. Guided by these data obtained in the single-celled eukaryote, *Tetrahymena thermophila*, experiments in vertebrate multiciliated cells revealed new insights into the mechanism of action for several poorly understood protein complexes associated with human ciliopathies. Together, this rich new resource and the experimental findings drawn from it highlight the power of evolutionary proteomics and structural biology to inform our understanding of human disease.

## Results and Discussion

### *In situ* interactome mapping of motile axonemes by cross-linking mass spectrometry

To map the protein interaction landscape of motile cilia (**Figure 1A**), we performed XL/MS on *Tetrahymena*, a single-celled ciliate that can be easily grown in massive quantities and whose ciliary beating machinery is highly conserved in vertebrates^37–39^. Cilia were isolated using standard protocols (see Methods and refs. ^33,39^) and incubated with the amine-reactive, membrane permeable, mass spectrometry-cleavable cross-linker, DSSO (disuccinimidyl sulfoxide). Cross-linked residues were identified by mass spectrometry and selected with a false discovery rate (FDR) of 1% and a Max XlinkX score > 40 (**Figure 1B**). This analysis yielded 19,339 cross-links, comprising 4,757 unique binary amino acid interactions measured for 1,143 proteins (**Figure 1C**). Of these, 1,377 XLs included two distinct proteins (intermolecular links), suggesting that they occurred at residue pairs lying within < 30 Å of one another. The remainder of XLs represented intramolecular interactions within proteins, providing information about the 3D structure of these proteins.

The machinery of motile axonemes is highly conserved across evolution^37^, and our ultimate goal was to leverage the accessibility of *Tetrahymena* cilia to improve our understanding of vertebrate motile cilia. We therefore interpreted our XL/MS data in the context of conserved eukaryotic protein families, specifically orthologs as defined using eggNOG^40^. We grouped the UniProt *Tetrahymena* proteome (see Methods and refs. ^41,42^) such that *Tetrahymena* proteins mapping to the same eukaryotic ortholog group (orthogroup) were considered together for purposes of interpretation and ready matching to their human orthologs. Thus, data throughout the paper will be presented using human protein names for ease of comparison, though in all cases, the UniProt names for the exact *Tetrahymena* proteins are provided in **Supp. Table 1**.

Figure 2A illustrates the breadth of the dataset, providing a protein-resolution visualization of just a small subset of the interactions identified among and between known machinery in motile cilia, including but not limited to outer dynein arms (ODAs), inner dynein arms (IDAs), the Nexin-dynein regulatory complex (N-DRC), radial spokes (RS), the tether/tether head complex, the central apparatus (CA) and numerous microtubule-associated proteins. Figure 2B illustrates the depth of the dataset, providing an amino acid-resolution visualization of the interactions identified among and between just five of the proteins from the network in 2A. This very small sampling revealed the potential impact of this dataset, so we next sought to validate the dataset by comparing our XL/MS interaction data to known ciliary protein interactions and structures.

**Figure 2:**
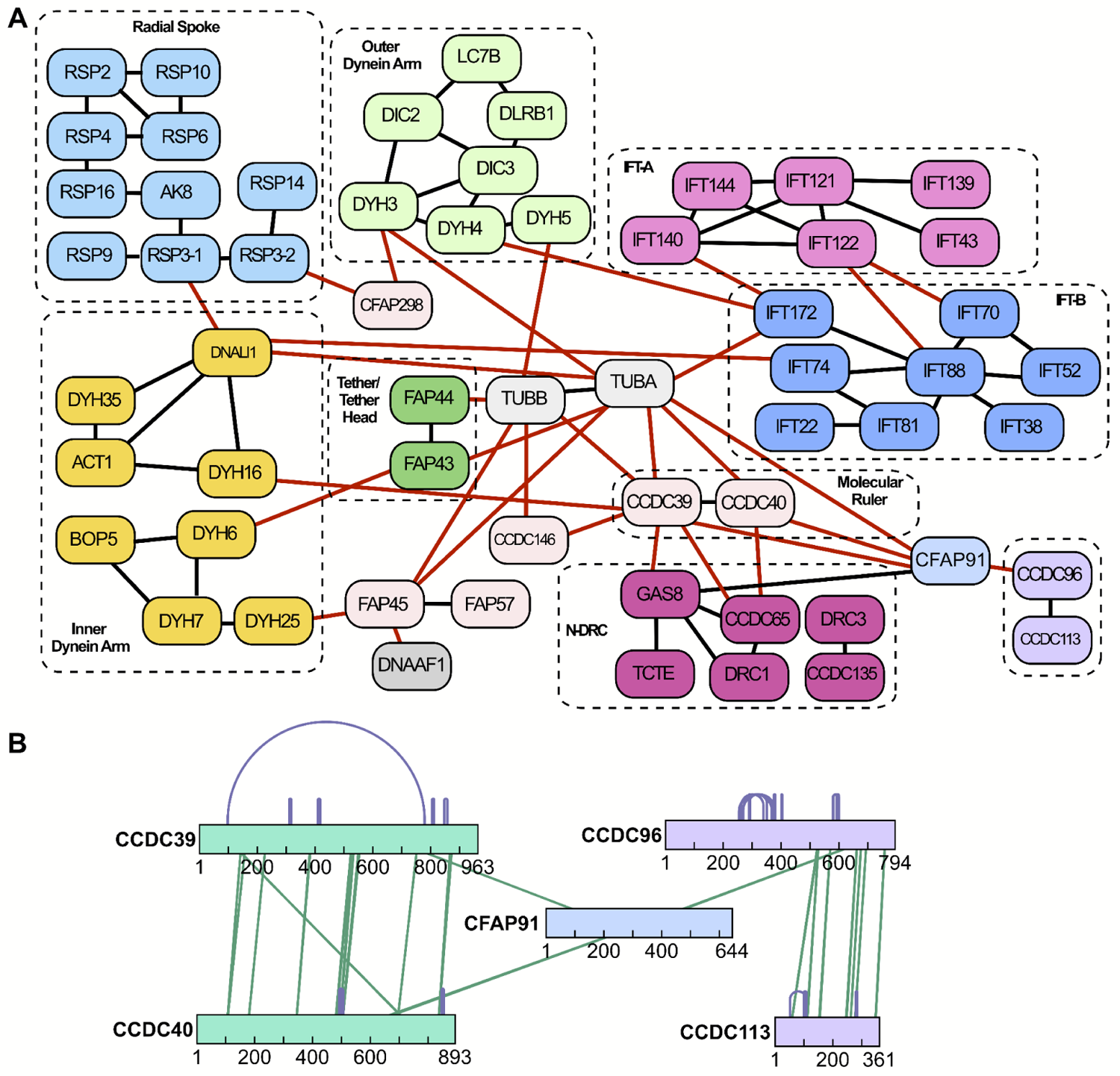
Cross-links highlight intra- and inter-complex interactions at amino acid resolution. (A) Binary protein interactions between proteins of the radial spoke (RS), outer dynein arm, inner dynein arm, IFT-A, IFT-B, tether/tether head, N-DRC, molecular ruler (MR), and several microtubule associated proteins. The binary protein interactions shown in red represent coverage of interactions between these protein complexes. While this diagram summarizes interactions at the protein level, the cross-links (XLs) capture more fine-grained (amino acid resolution) interactions, as illustrated by an expanded view of the CCDC96/CCDC113 complex and its partners in (B). Here, bars denote proteins labeled by amino acid positions, purple arcs indicate intramolecular XLs, and green lines indicate intermolecular XLs.

### XL/MS provides broad and deep coverage of the motile axoneme interactome

As an initial test of the validity of our XL/MS dataset, we mapped XLs onto several recently determined atomic-level axonemal structures and measured the distance between cross-linked atoms. Cross-links consistent with the existing models should be less than or equal to the 30 Å length (Cα-Cα) of the DSSO cross-linker and side chains^43^. Satisfyingly, when mapped to structures for *Tetrahymena* MIPs^44^ and *Chlamydomonas* radial spokes^45^, > 83% of 293 XLs across 30 proteins were found to measure within the expected 30 Å (**Supp.** Figure 1) and an additional 30 XLs (> 88%) measure within 40 Å. Intermolecular XLs tended, on average, to be longer than intramolecular XLs (**Supp.** Figure 1).

For a more granular test, we examined the XLs within the well-characterized IFT-A and IFT-B complexes^46,47^. Consistent with biochemical data^48,49^, we identified extensive XLs within each of the IFT-A and IFT-B complexes, but observed far fewer links between the two (Figure 3A).

**Figure 3:**
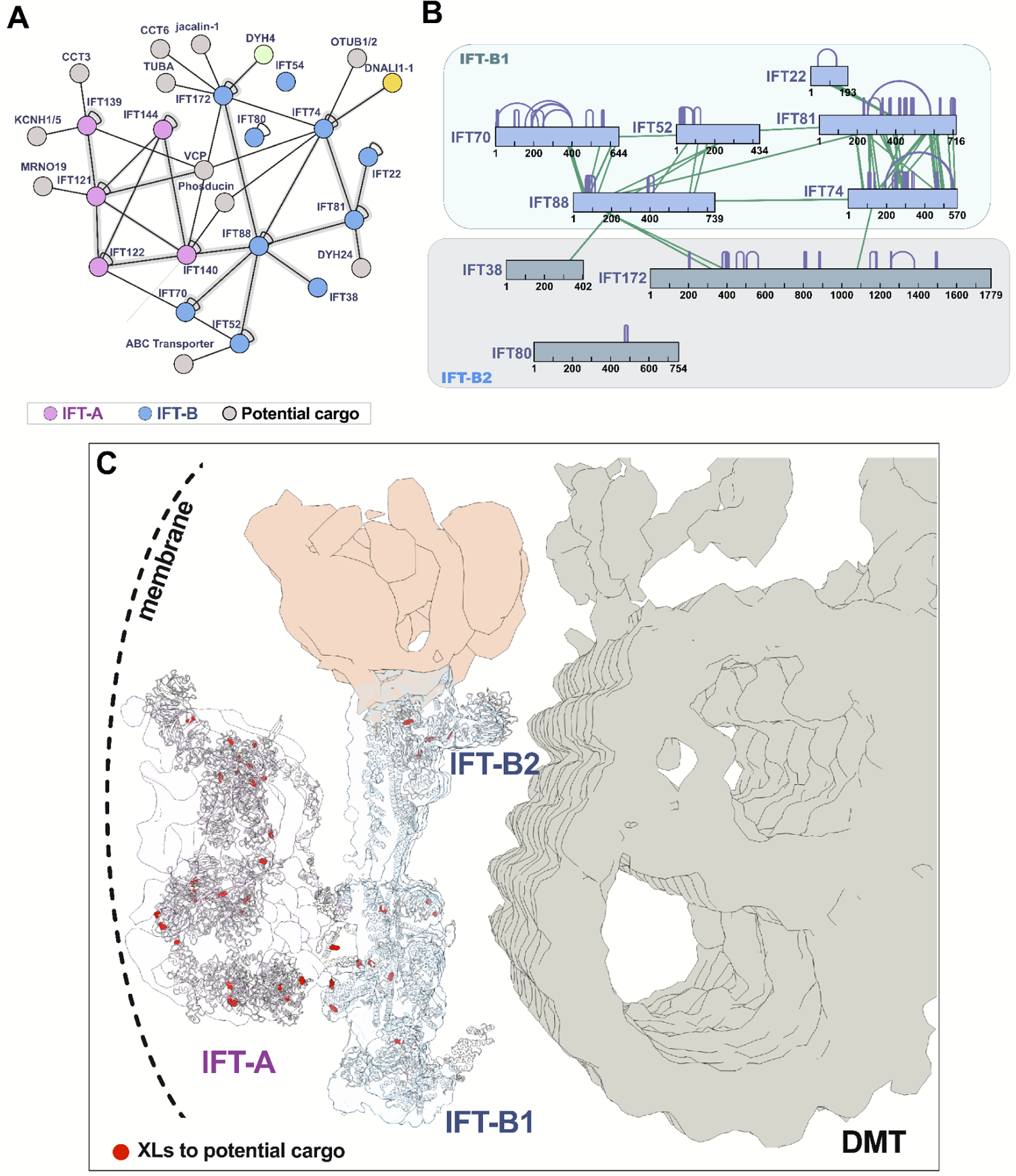
The interaction network of IFT proteins reveals stable structural interactions and transient associations with potential cargo. (A) The XL/MS data capture interactions between IFT-A proteins IFT140 and IFT122 to IFT-B proteins IFT88, IFT70, and IFT172, in addition to interactions to several potential IFT cargo proteins including dyneins and CCT chaperonin subunits. Fuzzy lines indicate ≥1 XL. (B) XLs within the IFT-B complex are enriched within the IFT-B1 and IFT-B2 subcomplexes with only sparse cross-links between them. (C) Molecular models of IFT-A and B (based on refs. ^33,100^) are superimposed onto the cryo-ET density from ref. ^101^, highlighting the generally membrane proximal positions of IFT-A amino acid residues (red circles) cross-linked to candidate cargo proteins.

Moreover, the observed cross-links precisely recapitulate recent cryo-electron tomography (cryo-ET) results, confirming that the two IFT complexes are linked by an intricate interaction network involving the IFT140 and IFT122 subunits of IFT-A and the IFT172, IFT88, and IFT70 subunits of IFT-B^50^ (Figures 3A**, B**). Likewise, IFT-B comprises two sub-complexes, IFT-B1 and B2^49,51–53^, and we again observed dense cross-linking within each of the sub-complexes. Finally, our XLs precisely reflected the 3.2 Å structure of IFT-B2^54^, with extensive interactions of IFT74 and IFT81 along their lengths, and a more discrete linkage to the N-terminus of IFT22 (Figure 3B). These comparisons demonstrate the high accuracy of our cross-linking data set.

The XL data from IFT complexes illustrate that the frequency of cross-links often reflects the stability of particular interactions but also suggest that the method can capture weak and/or transient interactions. This is important, because while protein-protein interactions within the IFT complexes are highly stable, the complexes’ role as transport machinery implies they must also more transiently interact with both regulatory modules and with cargos; such transient interactions may be more elusive^55^.

Excitingly, we clearly captured XLs between IFT proteins and known examples of both regulatory modules and cargoes. For example, the AAA ATPase Vcp interacts with and regulates IFT^56^, and our XL/MS identified direct association of this protein with both IFT-A and IFT-B (Figure 3A). Likewise,axonemal dyneins and radial spokes are known IFT cargoes^57–59^, and our data revealed links between IFT-B and both inner and outer axonemal dynein subunits and IFT-B and links between IFT-A and radial spoke proteins (Figure 3A, yellow, green, and light blue nodes, respectively). Moreover, IFT-A is specifically involved in entry of membrane protein to the cilium, and we found XLs directly between IFT-A and the ortholog of the cilia-related membrane channel Kcnh1^60^.

Finally, our data also identified new potential regulators or cargoes of IFT. For example, we observed XLs directly to both the CCT chaperonin and the phosducin PHLP2 (Figure 3A), both of which are essential for ciliogenesis *via* still-unknown mechanisms^61–63^. Interestingly, IFT residues participating in XLs with potential cargo proteins tended to form distinct clusters on the outer surfaces of IFT-A and IFT-B (Figure 3C).

### XL/MS captures conformational dynamics of motile ciliary protein complexes

Next, we reasoned that because our experiment included *in situ* cross-links, the XL/MS data should capture the dynamics of multi-protein machines. To test this idea, we examined the dozens of XLs linking subunits of the axonemal dyneins (Figure 4A). The dynamic nature of these massive motor complexes has been captured by three different atomic structures representing distinct phases of the motor’s duty cycle (Protein Data Bank (PDB) IDs 7K58, 7K58, 7KEK)^64^. We mapped our XLs onto these structures and measured the distance between cross-linked residues. We found clear concordance for most XLs, however, the dynamics of these motors was most clearly revealed by XLs that violated the predicted 30 Å length constraint, including many extreme violations (> 100 Å)(Figure 4B).

**Figure 4:**
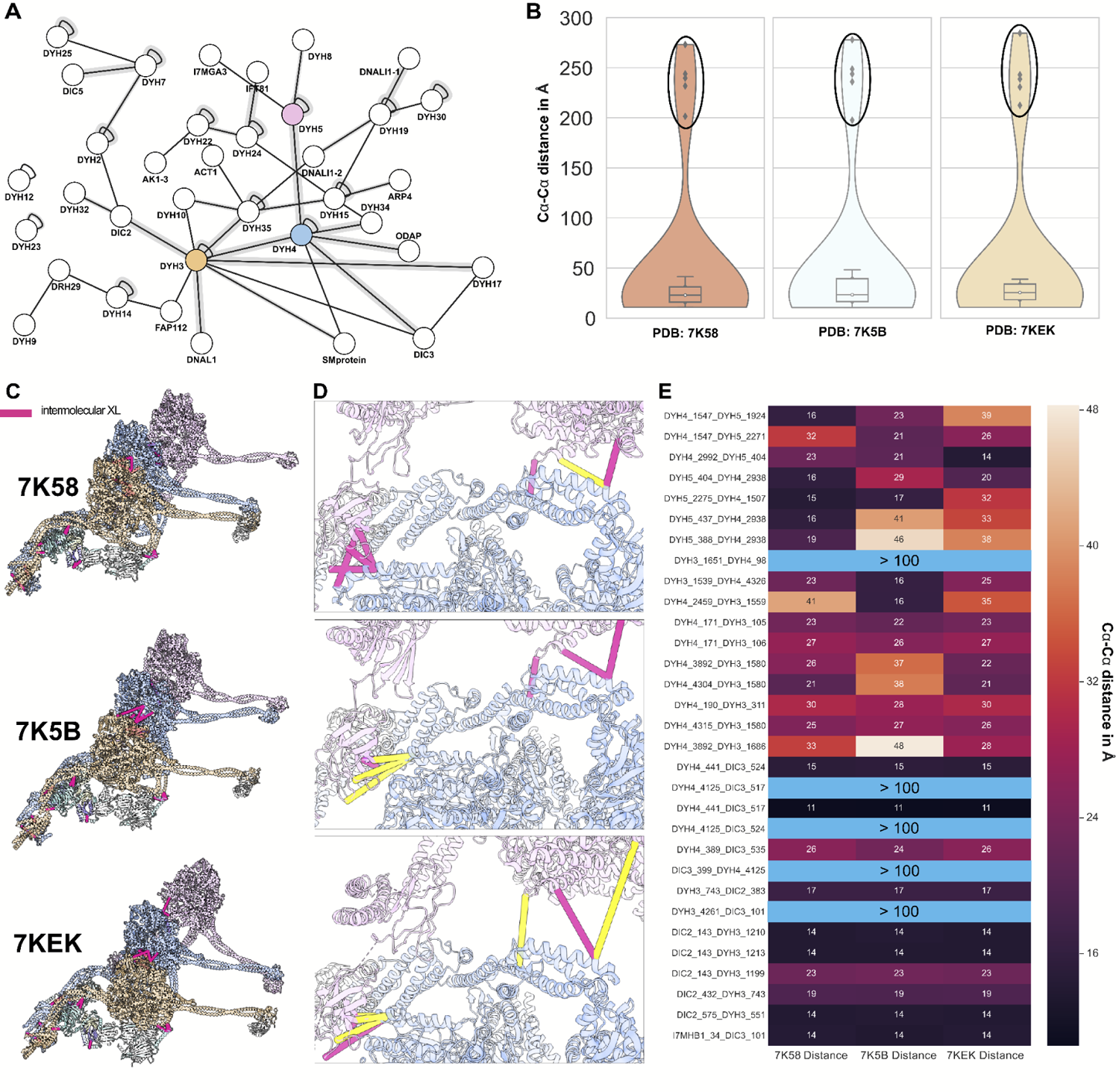
Cross-link distances emphasize dynamics of the outer dynein arm. (A) XL/MS reveals a network of interactions of outer dynein arms. When mapped onto the structure of the ODA (PDBIDs 7K58, 7K5B, 7KEK), intermolecular XLs stitch the components of the ODA proteins together, shown by (B) distance distributions and (C) as plotted onto the structures. (D) Close-ups of XLs show the dynamic shift of the complex between binding state 1 (top panel), binding state 2 (middle panel), and the pre-parallel state (bottom panel) with yellow showing violated XLs and pink satisfied XLs. (E) The heatmap of calculated XL distances from the ODA in different conformations shows that all XLs are satisfied by at least one of the conformations. The large intramolecular violations circled in (B) and highlighted in (E) in light blue are satisfied except for one (slight violation of 37 Å) by intermolecular interactions within the *in situ* head-to-tail conformation of the ODA.

We reasoned that the moderate violations in any given structure may reflect XLs in complexes with a different conformation, so we mapped our XLs to the different structures (Figure 4C).

Indeed, nearly all length violations for any single conformation (Figure 4D**, yellow**) could be satisfied when mapped to another of the conformations (Figure 4D**, pink**). The heatmap in Figure 4E reveals that all observed violations could be satisfied by mapping to at least one of the multiple conformations, with the exception of the extreme (> 100 Å) violations.

Crucially, we were able to satisfy these extreme violations by taking into account the oligomerization of multiple ODAs, previously described to be present in a tail-to-head arrangement along the axoneme^64^. When interpreted as inter-links between residues on adjacent motor complexes, these XLs now were < 30Å.

Thus, while additional positive controls will be presented throughout the paper, these initial tests strongly suggest that our XL/MS dataset provides broad, dynamic coverage of the ciliary beating machinery, captures both relatively stable and relatively transient interactions, and accurately informs not only individual protein-protein interactions, but also protein structures and protein complex conformations in the motile axoneme. We therefore used our interactome map to interrogate the functions of several poorly defined protein complexes related to human motile ciliopathy.

### A diverse adenylate kinase repertoire in the central apparatus and radial spokes of motile axonemes

Adenylate kinases that catalyze interconversions of adenosine phosphates are enriched in energy-intensive organelles such as motile cilia, including in *Tetrahymena*^65,66^. Among these, AK8 has been found to interact with the human ciliopathy protein CFAP45^67^, but while work in mice demonstrates AK8 is required for cilia beating^68^, its position in the axoneme and its mechanism of action remain unclear. For example, the interaction with CFAP45 suggests proximity to MIPs inside the doublet microtubules (DMT), yet AK8 is missing in axonemes lacking the RS1 radial spoke, which lies outside the DMTs^69^. We were interested then to find several XLs in *Tetrahymena* linking AK8 to the RS1 radial spoke protein RSPH3, but not to MIPs (Figure 5A). To gain further insight, we examined a recent cryo-EM map of the *Chlamydomonas* radial spokes, and we identified an unassigned protein backbone in RS1^70^. Strikingly, we found that an AlphaFold-predicted dimer of AK8 fit very well within the fold of the unassigned protein, in addition to satisfying the XL between AK8 and RSPH3 (RMSD of 2.3 Å when superimposed onto an unassigned protein in PDBID 7JTK)(Figure 5B). These data suggest a role for AK8 in the *Tetrahymena* RS1 radial spoke.

**Figure 5:**
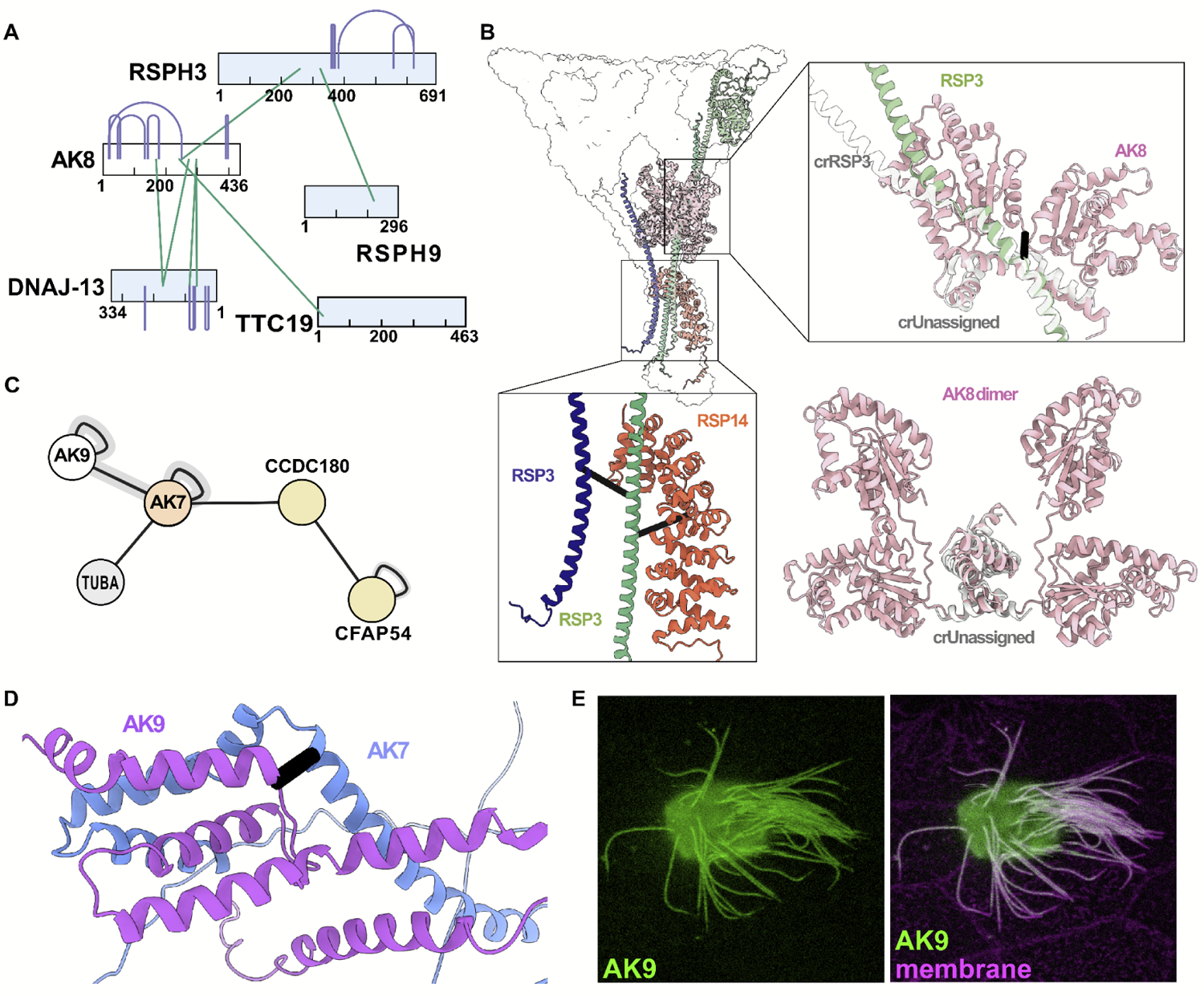
Adenylate kinases participate in several major ciliary complexes. (A) Cross-links capture interactions between AK8 and radial spoke proteins. **(B)** The structure of the *C. reinhardtii* radial spoke 1 (PDBID 7JTK) shows an unassigned protein with the same fold as the AK8 dimer predicted by AlphaFold2. The AlphaFold2 AK8 dimer and unassigned protein can be superimposed with an RMSD of ∼2 Å; moreover, all AK8 intramolecular XLs and AK8-RSP3 intermolecular cross-links are satisfied by this assignment. **(C)** Cross-links show an association between AK7 and AK9 agreeing with **(D)** the AlphaFold2 model of the C-terminus of the two proteins (black bar denotes XL position). AK7 is also cross-linked to central apparatus protein CCDC180/CFAP76. **(E)** An *en face* view of the apical surface of a *Xenopus* multiciliated cell reveals that AK9 is present in ciliary axonemes, right panel shows counter-staining with membrane-RFP. Scale bar = 10 µm.

Another ciliary adenylate kinase, AK7 is essential for ciliary beating and is linked to human motile ciliopathies associated with defects in the central apparatus^71,72^, though little else is known of this protein’s function or site of action. We observed several XLs linking the AK7 ortholog in *Tetrahymena* to both tubulin and to CCDC180/CFAP76 (Figure 5C), a component of the C1c structure of the central apparatus^73,74^. Moreover, we observed several XLs linking AK7 to AK9, and with AlphaFold2, we modeled the region of the interaction between the proteins (Figure 5D). This result is of interest because while the enzymatic activity of vertebrate AK9 has been defined *in vitro*^75^, nothing is known of this protein’s localization or function *in vivo*. This result prompted us to examine a GFP fusion to AK9 expressed in *Xenopus* epidermal multiciliated cells (MCCs); we found that this protein robustly localized along the length of the axoneme (Figure 5E). Our data thus argue that AK7 and AK9 likely form a heterodimer and function together as integral components of central apparatus of vertebrate motile axonemes. Our XL/MS dataset thus provides novel insights into the action of several ill-defined adenylate kinases in motile cilia.

### A dense network of ciliopathy proteins associated with the CCDC39/40 96nm ruler

CCDC39 and CCDC40 form the so-called “96nm ruler” that sets the repeat length of the axonemal dyneins, and disruption of the complex results in severe derangement of the axoneme and especially poor patient outcomes^9–11^. We therefore sought to better define the protein interaction landscape around CCDC39/40. As a positive control, we noted that our XLs clearly reflect the existing ∼6 Å cryo-ET structure of the complex^12,70^, showing the intimate association of CCDC39 and CCDC40 along their lengths as well as their interaction with the nexin-dynein regulatory complex (N-DRC) and the doublet microtubules (Figure 6A). Excitingly, our XL/MS also revealed several novel interactions (Figure 6B).

**Figure 6:**
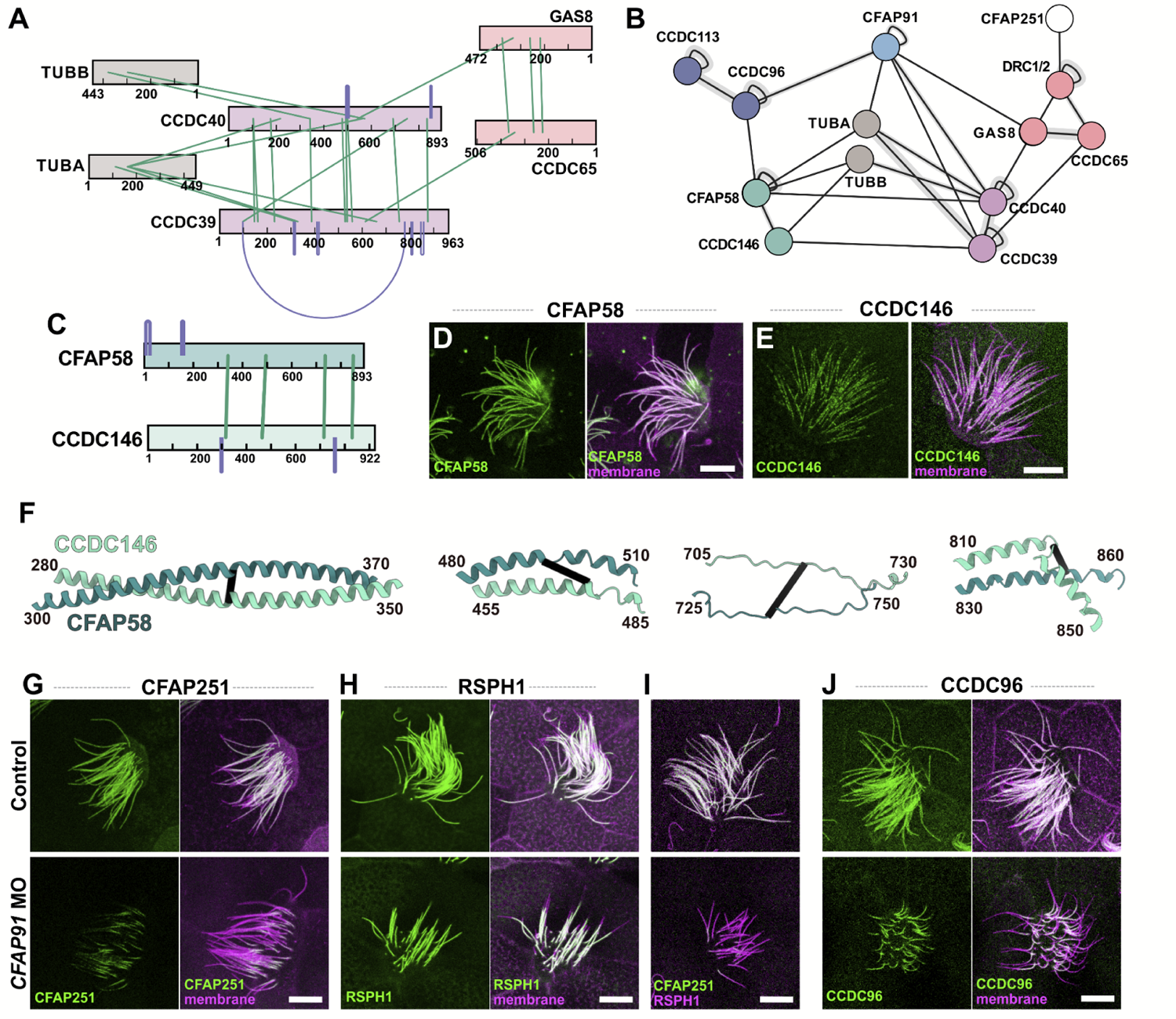
The molecular ruler (CCDC39/40) is a hub for proteins implicated in ciliopathies. (A) The molecular ruler forms dense interactions between proteins CCDC39 and CCDC40 in addition to interactions with the DMT and proteins of the N-DRC complex captured by our XL/MS. (B) The broader interaction network from our data shows the molecular ruler interacting with poorly understood ciliopathy protein and RS member, CFAP91. The molecular ruler also interacts with poorly classified DMT associated proteins CFAP58 and CCDC146. (C) Cross-links within and between CFAP58 and CCDC146 suggest a coil-coil is formed in a head-to-head conformation of the two proteins. (D)-(E) Confocal live images visualized by membrane-RFP and GFP-CFAP58 (D) or GFP-CCDC146 (E). Both CFAP58 and CCDC146 localize to the axonemes in *Xenopus* multiciliated epithelial cells. (F) Selected chunks of the AlphaFold2 predicted dimer between CFAP58 and CCDC146 are in agreement with the cross-link from the motile cilia proteome. (G)-(H) After *CFAP91* knockdown (Supp. Figure 2), GFP-CFAP251 (G) failed to localize to the axoneme, while GFP-RSPH1 (H) remained unaffected. (I) The specific CFAP251 loss was confirmed by dual labeling of mCherry-RSPH1. (J) CCDC96 failed to localize to the axoneme after *CFAP91* knockdown. Scale bars = 10 µm.

For example, CCDC39 and CCDC40 interacted, respectively, with CCDC146 and CFAP58 (Figure 6B**, teal**). Both latter proteins are essential for cilia beating but their function and even their position within the axoneme are unknown^76–78^. Interestingly, our XL/MS data revealed extensive interaction along the two lengths of CCDC146 and CFAP58, suggesting a stable complex (Figure 6C**, 6F**), and XLs linking the complex to the exterior of tubulin suggest the complex lies outside the DMT. We found as well that both proteins localize to motile cilia in *Xenopus* MCCs (Figure 6D**, E**).

We also identified multiple XLs connecting CCDC39/40 and N-DRC subunits to CFAP91/MAATS1(Figure 6B**, light blue**), which is essential for cilia beating and is associated with human motile ciliopathy but is otherwise poorly understood^79–81^. A recent EM study showed that this elongated protein forms the base of the RS2 radial spoke, extends parallel to the CCDC39/40 ruler and through the N-DRC, and contacts RS3^70,82^. Moreover, defective CFAP91 function in both humans and *Tetrahymena* is associated with disruption specifically of the RS3 radial spoke^79,81^. Little else is known of CFAP91 function, so it is significant that multiple XLs in our dataset also linked CFAP91 directly to CCDC96 (Figure 6B**, dark blue**), which connects RS3 to the NDR-C^83^.

These interactions in *Tetrahymena* prompted us to explore the effect of CFAP91 loss in vertebrate multiciliated cells. First, we showed that knockdown (KD) of CFAP91 in *Xenopus* multiciliated cells recapitulated the known effect of genetic loss in humans and *Tetrahymena*^79,81^. For example, we observed specific loss of the RS3 subunit CFAP251 (Figure 6G), while the RS1/RS2 subunit Rsph1 was unaffected (Figure 6H). We confirmed the specificity of CFAP251 loss by double labeling with RSPH1(Figure 6I). We then used this manipulation to test and explore the significance of a novel, direct interaction identified by our XL/MS between CFAP91 and CCDC96 (Figure 6B, blue); CFAP91 KD elicited a striking loss of CCDC96 from axonemes (Figure 6J). Thus, discovery-based proteomics in *Tetrahymena* provided an amino-acid level view of the intimate network formed by CFAP91 and components of RS2, RS3, the N-DRC, and the 96nm ruler that enabled targeted experiments in *Xenopus* to elucidate the specific and complex molecular phenotype arising from this ciliopathy network’s disruption.

### XL/MS-directed analysis of links between MIPs and axonemal dyneins illuminate the etiology of poorly understood motile ciliopathy proteins

The MIPs form complex networks lining the inner surface of microtubule doublets in motile axonemes and play a role in stabilizing their structure^84,85^. In addition, some MIPs penetrate the MT wall to contact and regulate the axonemal dyneins outside^86^. Previously, we and others showed that disruption of one MIP, ENKUR, was associated with human ciliopathy and is required for sperm flagellar beating and normal left/right patterning, yet no known structural or molecular phenotype has yet been found in ENKUR deficient axonemes^87,88^. We therefore used our XL/MS data to guide a deeper investigation of this protein’s function.

ENKUR is a MIP and is known to be positioned within the B tubule of the MT doublet^44^, and accordingly, our XL/MS data linked ENKUR directly to another B-tubule MIP, CFAP45, but not to other nearby MIPs, such as CFAP52 or PACRG (Figure 7A). Consistent with our interactome data, knockdown of ENKUR elicited a robust loss of CFAP45 localization in motile axonemes in *Xenopus* MCCs, providing the first known defect in ENKUR deficient axonemes. The effect was highly specific, as CFAP52 was largely unaffected by ENKUR KD (Figure 7C, D), a result confirmed by dual-labeling for CFAP45 and CFAP52 (Figure 7E).

**Figure 7:**
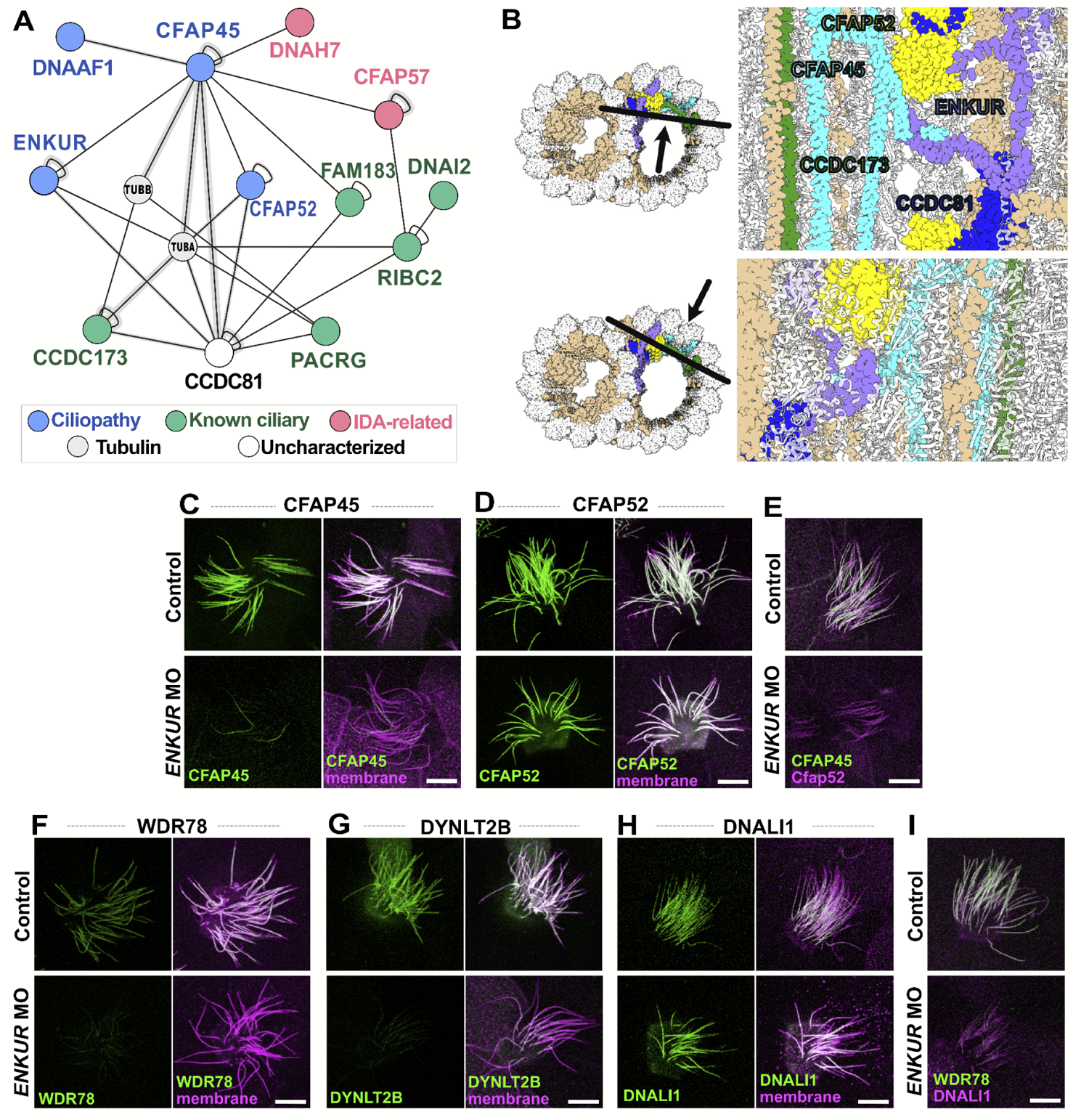
XL/MS helps to reveal the intricate interaction network of MIPs CFAP45 and ENKUR along the DMT. (A) Cross-linking network surrounding CFAP45 and ENKUR showing a network of ciliopathy-related proteins. (B) The molecular structure of the native *Tetrahymena* microtubule doublet confirms interactions shown in (A). (C)-(D) After *ENKUR* knockdown, GFP-CFAP45 (C) failed to localize to the axoneme, while GFP-CFAP52 (D) remained unaffected. (J) The specific CFAP45 loss was confirmed by dual labeling of mCherry-CFAP52. (F)-(H) After *ENKUR* knockdown, IDA(f) subunits, GFP-WDR78 (F) and GFP-DYNLT2B (G) failed to localize to the axoneme, while GFP-DNALI1 (H), an IDA(a, c, d) subunit remained unaffected. (I) The specificity of IDA(f) was confirmed by dual labeling of mCherry-DNALI1. Scale bars = 10 µm.

This result, while consistent with our XL/MS data, does not explain the ciliary beating defect in ENKUR deficient cilia, as CFAP45-deficient cilia also appear largely normal^67^. We therefore explored deeper into the ENKUR/CFAP45 protein interaction neighborhood. Our XL/MS demonstrate that the association involves the N-terminus of CFAP45 and C-terminus of ENKUR, regions otherwise unresolved in the recently published structure of the native *Tetrahymena* doublet microtubule (DMT)^89^. Moreover, the structure raises the possibility that like other MIPs^86^, ENKUR and/or CFAP45 may penetrate the MT doublet wall. Indeed, this interpretation is strongly supported by XLs connecting CFAP45 to two IDA-related proteins, the dynein heavy chain DNAH7 and the IDA adaptor, CFAP57 (Figure 7A**, pink**). CFAP57 was of particular interest, as it is linked to motile ciliopathy in humans^13^ and its loss in *Chlamydomonas* elicits specific defects in the assembly of only a single, specific subtype of IDAs, IDA(f)^90^. These findings prompted us to test the idea that ENKUR loss may specifically impact IDA(f) assembly.

Indeed, ENKUR loss in *Xenopus* severely disrupted the axonemal deployment of two distinct IDA(f) subunits, WDR78 and DYNLT2B (Figure 7F**, G**). This effect was highly specific, as the IDA(a,c,d) subunit DNALI1 is normally deployed in mouse mutants of ENKUR^87^, and we found here that DNALI1 is also retained in *Xenopus* MCC axonemes after Enkur loss (Figure 7H**; also see Supp.** Figure 3). The specificity of the IDA(f) defect in *Xenopus* was confirmed by dual labeling of WDR78 and DNALI1 (Figure 7I). Thus, our XL/MS data from *Tetrahymena* guided the discovery of the first known molecular defect resulting from loss of the ciliopathy protein ENKUR.

## Conclusions

Motile cilia occupy a central position in both fundamental biology and biomedicine. Present in the last eukaryotic common ancestor, these organelles play essential roles in organisms across the tree of life^38^. Not only do they drive locomotion and sensation for organisms in aqueous environments, but they also generate fluid flows essential for the development and homeostasis of several different organ systems in more complex animals^91–93^. Accordingly, defects in the structure and function of motile cilia underlie a wide spectrum of human genetic diseases, the motile ciliopathies^1,2^. Here, we exploited the strong evolutionary conservation of motile cilia to provide insights into the composition of ciliopathy-associated protein complexes. Taking advantage of the hundreds of cilia decorating the surface of *Tetrahymena*, we generated a motile cilia interactome of 9,208 unique ciliary protein interactions covering essentially all elements of the motile cilia machinery.

Importantly, we identified several novel interactions for poorly defined ciliopathy proteins. For example, we shed light on the essential role of adenylate kinases in the cilia through their interactions with radial spoke proteins and the central apparatus. In addition, we highlight two new microtubule-associated proteins, CFAP58 and CCDC146, both of which are implicated in ciliary dysfunction. We show these proteins form a coil-coil and map to the exterior of the DMT, forming interactions with the molecular ruler (CCDC39/40), CCDC96/113, and Rsp3. Most importantly, our proteomic analyses led us to discover the first molecular defects in cilia lacking the ciliopathy protein ENKUR, revealing a specific defect in the localization of IDA(f) to the ciliary axoneme.

This proteomic resource, therefore, should be of broad utility, informing both the basic biology of cilia and the etiology of ciliopathies. For example, genetic diagnosis of motile ciliopathy is a complex endeavor, and the long-standing method of TEM analysis of cilia ultrastructure is insufficient in many cases to identify functional defects^15^. Recently, new diagnostic methods have been developed that exploit the power of super-resolution imaging and the localization of specific proteins in motile cilia^94^. Accordingly, better definition of not only normal protein localization in axonemes but also of proteins’ functional interactions will be critical.

Finally, the depth and breadth of the protein-protein interactome generated here highlight the power of XL/MS as a transformative tool in uncovering intricate molecular networks associated with cellular processes. We demonstrate the value of evolutionary conservation in bridging proteomics and *in vivo* studies to advance our understanding of complex systems and their implication in human disease. The knowledge derived from this study provides valuable insights into the molecular basis of ciliopathies and suggests new aspects and potential targets for future therapeutic interventions.

## Materials and Methods

### Tetrahymena thermophila culture and deciliation

*T. thermophila* SB715 obtained from the Tetrahymena Stock Center (Cornell University, Ithaca, NY) were grown in Modified Neff medium at ∼ 21°C. Cilia were isolated from freshly grown 3-4 Liter cultures grown with shaking (100 rpm) by pH shock or dibucaine treatment using Hepes-buffered Cilia Wash Buffer as in McCafferty *et al*.^43^. Membrane and Matrix (M+M) extract was made by addition of 1% NP40 as in ref. ^33^.

#### Protein cross-linking

Cross-linking was performed as previously described^43^. Briefly, freshly dissolved DSSO (50 mM in DMSO or DMF) was added to either intact cilia isolated as above or to native chromatographic fractions of M+M extract separated by non-denaturing chromatography, using either SEC or mixed bed IEX. Cross-linking reactions were incubated for 60 minutes at room temperature (∼21°C) then quenched by addition of Tris pH 8.0 to > 20 mM. Proteins were solubilized with SDS and precipitated with acetone before trypsin digestion and desalting. Cross-linked peptides were enriched by chromatography on a GE Superdex 30 Increase 3.2/300 SEC column with approximately 6 cross-link-containing fractions dried and resuspended for mass spectrometry.

#### MS^3^ mass spectrometry and identification of cross-links

Data were collected as previously described^43^. Briefly, all data were collected on a Thermo Orbitrap Fusion Lumos tribrid mass spectrometer. A standard DDA LC/MS-MS workflow was used to identify proteins present and generate a *.fasta* file for cross-link identification. Spectra for cross-link analysis were collected using 115 min or 185 min gradient DDA MS2-MS3 methods. Mass spectrometry data were analyzed as follows: Lys-Lys cross-links were identified using the standard DSSO cleavable cross-link workflow in the XlinkX node of Proteome Discoverer 2.3 (Thermo) as described in ref.^32^ and results exported to xiView^95^ for visualization.

#### AlphaFold2 structure prediction

The protein structure predictions presented in the manuscript were predicted using the 2.2 release of AlphaFold2^96^ multimer implemented on the Texas Advanced Computing Center (TACC) cluster^97^.

#### Manipulation of Xenopus embryos

Female adult *Xenopus laevis* were induced to ovulate by injection of hCG(human chorionic gonadotropin). *In vitro* fertilization was carried out by homogenizing a small fraction of a testis in 1X Marc’s Modified Ringer’s (MMR). Embryos were dejellied in 1/3X Marc’s Modified Ringer’s (MMR) with 2.5%(W/V) cysteine at pH 7.8 and were microinjected with mRNA or morpholino in 2% Ficoll (W/V) in 1/3X MMR at the 4-cell stage. Injected embryos were washed with 1/3X MMR after 30 min and were incubated until tadpole stages.

*Xenopus* gene sequences were obtained from Xenbase (www.xenbase.org). Gene open reading frames (ORF) were amplified from the *Xenopus* cDNA library by polymerase chain reaction (PCR), and then are inserted into a pCS10R MCC vector containing a fluorescence tag.

Capped mRNAs were synthesized using mMESSAGE mMACHINE SP6 transcription kit (ThermoFisher Scientific). Morpholinos (MO) were designed to block splicing (Gene Tools). The *ENKUR* MO sequence was 5’-AATGACTATCCACTTACTTTCAGCC-3’ ^87^ and *CFAP91* MO sequence was 5’-AAATGGAGGCAGTACAGGTACATAC-3’. 40∼80 pg of each mRNA or 20ng of each MOs was injected into two ventral blastomeres.

*Xenopus* embryos were mounted between cover glass and submerged in 1/3x MMR at tadpole stages, and then were imaged immediately. Live images were captured with a Zeiss LSM700 laser scanning confocal microscope using a plan-apochromat 63X/1.4 NA oil objective lens (Zeiss) or with Nikon eclipse Ti confocal microscope with a 63×/1.4 oil immersion objective. Captured images were processed in Fiji.

To verify the efficiency of *CFAP91* MO, MO was injected into all cells at the 4-cell stage and total RNA was isolated using the TRIZOL reagent (Invitrogen) at stage 25. cDNA was synthesized using M-MLV Reverse Transcriptase (Invitrogen) and random hexamers (NEB). CFAP91 cDNA was amplified by Taq polymerase (NEB) with the following primers:

*CFAP91* 65F 5’-GAGCGTACGACTTCCTCTATGA-3’

*CFAP91* 541R 5’-GTCACGGTAATCCGTCTGAATG-3’

#### Data deposition

All mass spectrometry proteomics data and their full reanalyses described in this paper are deposited in the MassIVE/ProteomeXchange databases (https://massive.ucsd.edu, see also ref. ^98^ under MassIVE/ProteomeXchange accession numbers MSV000089917 / PXD035387 (*in situ* cilia cross linking dataset, denoted XL1), MSV000090056 / PXD035702 (*in situ* cross-linking then solubilized dataset, denoted XL2), and MSV000089131 / PXD032818 (cross-linking of partially purified IFTA (FPLC SEC) and cross-linked cilia extract IEX fractions). Pairwise protein interactions were deposited in the IntAct database^99^ and are additionally available, along with other supporting materials, including 3D models, in a supporting Zenodo repository available at 10.5281/zenodo.8360038.

## Acknowledgements

The authors gratefully acknowledge the generous support of the *Tetrahymena* stock center (Cornell University & Washington University in St. Louis). Research was funded by grants from the National Institute of General Medical Sciences R35GM122480 (to E.M.M.) and R35GM138348 (to D.W.T.), National Science Foundation (2019238253 to C.L.M.), National Institute of Child Health and Human Development (HD085901 to J.B.W. and E.M.M.), Army Research Office (W911NF-12-1-0390 to E.M.M.), and Welch Foundation (F-1515 to E.M.M., F-1938 to D.W.T.). D.W.T. is a CPRIT Scholar supported by Cancer Prevention and Research Institute of Texas (RR160088). The authors acknowledge the Texas Advanced Computing Center at The University of Texas at Austin for providing high-performance computing resources that have contributed to the research results reported within this paper.

## Competing interests

The authors declare no competing interests. E.M.M. is a co-founder, shareholder, and scientific advisory board member of Erisyon, Inc., which played no role in this work.

## Supplementary Figures

**Supplementary Figure 1.**
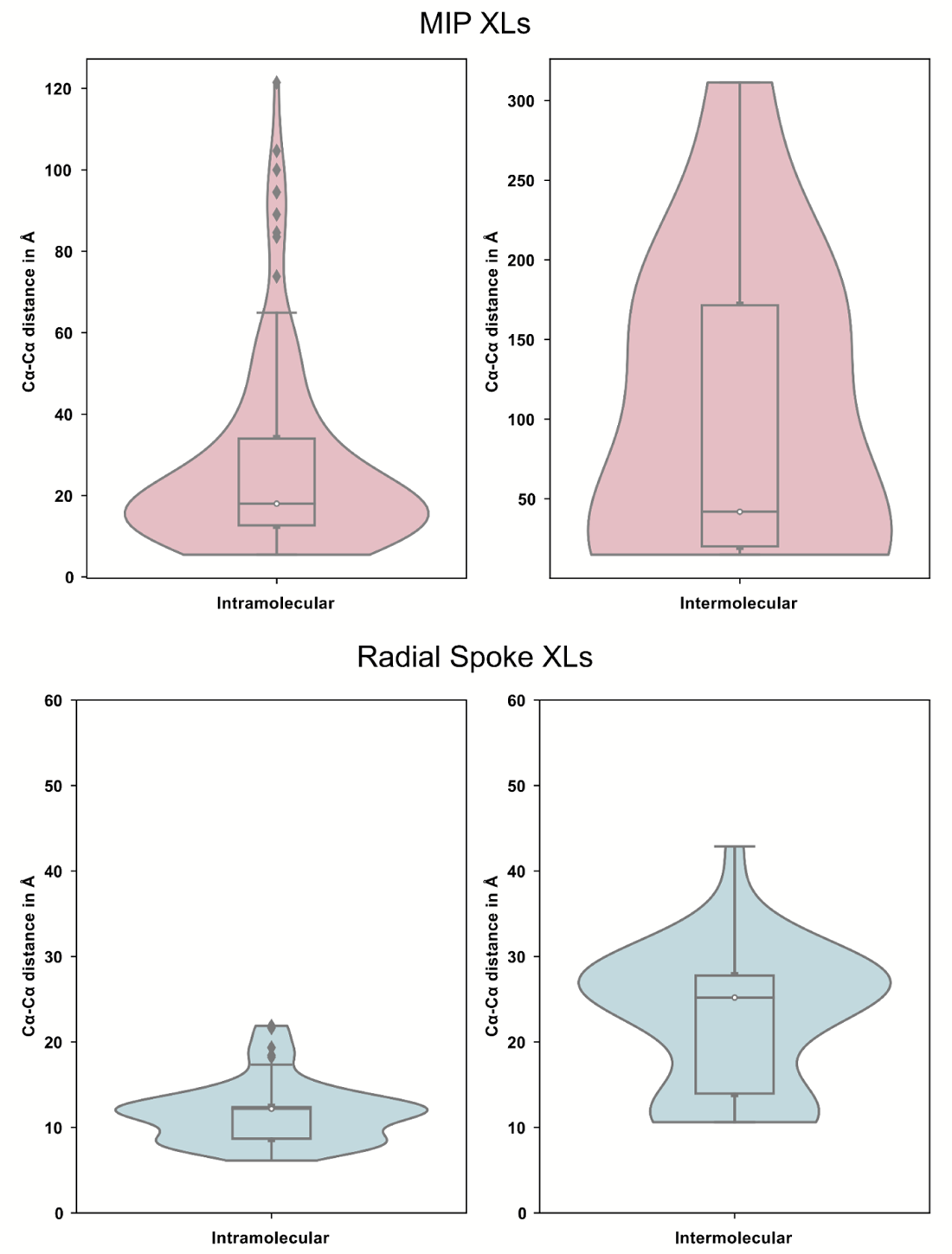
Distance distribution of intra- and intermolecular cross-links of ciliary MIP and radial spoke proteins.

**Supplementary Figure 2.**
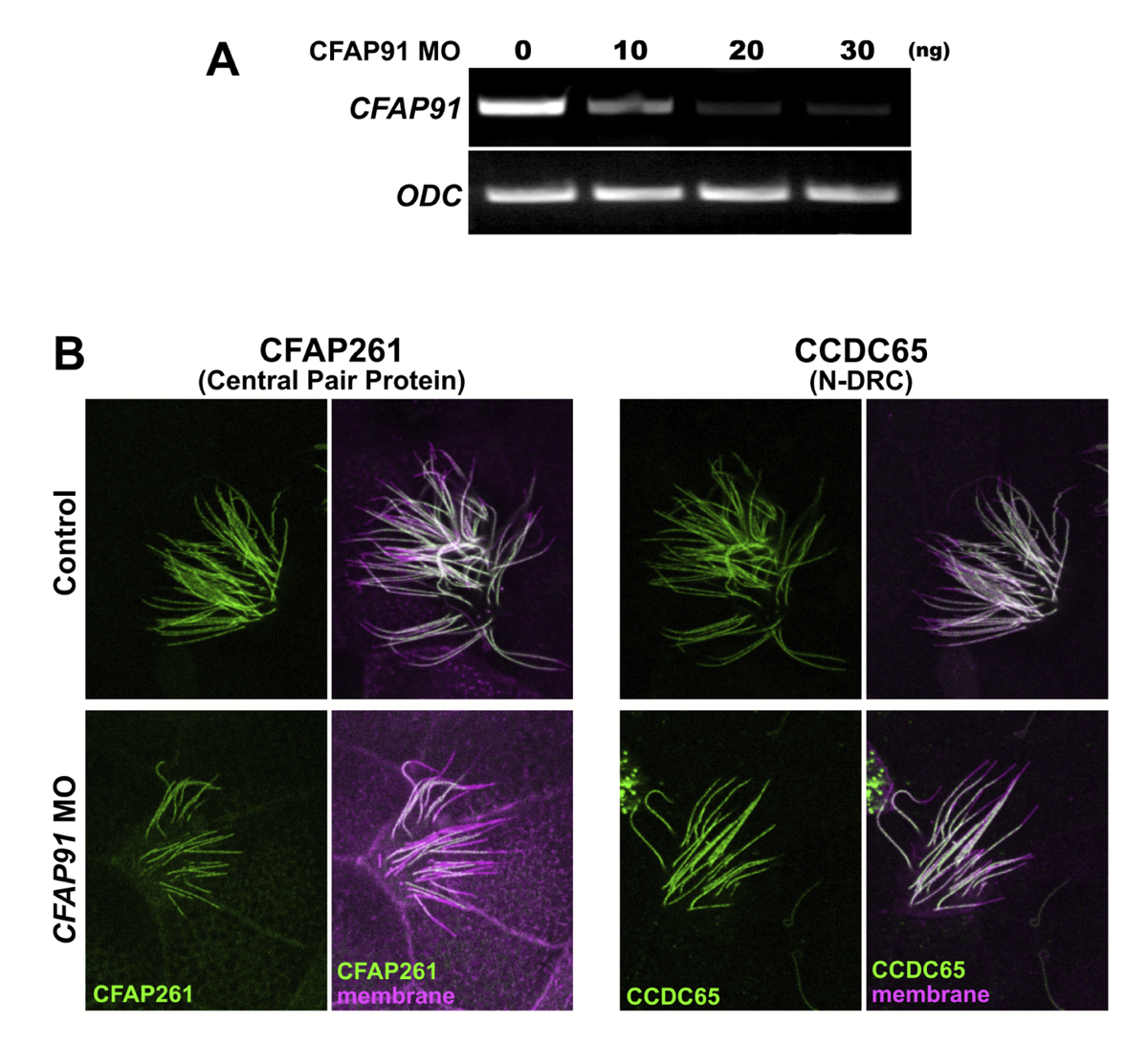
(A) RT-PCR showed effective disruption of splicing by injection of *CFAP91* morpholino. (B) Localization of GFP fusions to indicated proteins in *Xenopus* MCCs.

**Supplementary Figure 3.**
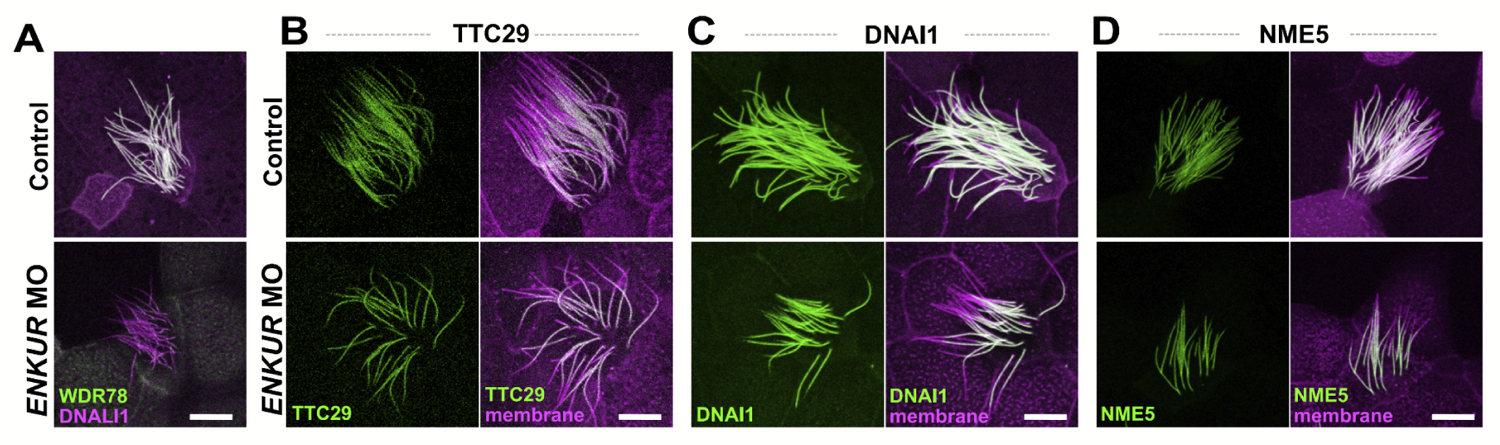
**(A)** After ENKUR knockdown, GFP-WDR78 (an IDA-f subunit) failed to localize to the axoneme, while mCherry-DNALI1 remained unaffected. **(B)-(D)** The loss of *ENKUR* did not affect the localization of TTC29 (an IDA-d subunit), DNAI1 (an ODA subunit) and NME5 (a radial spoke protein) on the axonemes. Scale bars = 10 µm

---

**Supplementary Table 1.** List of unique cross-links with Tetrahymena and Uniprot gene identifiers, cross-linked amino acid positions, EggNOG (v. 5) orthogroup identifiers, and human orthologs.

## Notes

### Competing Interest Statement

The authors have declared no competing interest.

### Summary of Updates

In light of new questions raised about the accuracy of non-lysine disuccinimidyl sulfoxide cross-links in cross-linking mass spectrometry, the new version reports a more conservative dataset relying on only lysine-lysine cross-links.

